# Data Independent Acquisition to Inform the Development of Targeted Proteomics Assays Using a Triple Quadrupole Mass Spectrometer

**DOI:** 10.1101/2024.05.29.596554

**Authors:** Deanna L. Plubell, Eric Huang, Sandra E. Spencer, Kathleen Poston, Thomas J. Montine, Michael J. MacCoss

## Abstract

Mass spectrometry based targeted proteomics methods provide sensitive and high-throughput analysis of selected proteins. To develop a targeted bottom-up proteomics assay, peptides must be evaluated as proxies for the measurement of a protein or proteoform in a biological matrix. Candidate peptide selection typically relies on predetermined biochemical properties, data from semi-stochastic sampling, or by empirical measurements. These strategies require extensive testing and method refinement due to the difficulties associated with prediction of peptide response in the biological matrix of interest. Gas-phase fractionated (GPF) narrow window data-independent acquisition (DIA) aids in the development of reproducible selected reaction monitoring (SRM) assays by providing matrix-specific information on peptide detectability and quantification by mass spectrometry. To demonstrate the suitability of DIA data for selecting peptide targets, we reimplement a portion of an existing assay to measure 98 Alzheimer’s disease proteins in cerebrospinal fluid (CSF). Peptides were selected from GPF-DIA based on signal intensity and reproducibility. The resulting SRM assay exhibits similar quantitative precision to published data, despite the inclusion of different peptides between the assays. This workflow enables development of new assays without additional up-front data acquisition, demonstrated here through generation of a separate assay for an unrelated set of proteins in CSF from the same dataset.

## INTRODUCTION

Mass spectrometry (MS)-based targeted proteomics is a powerful alternative to immunoassays for the precise and accurate quantification of proteins in complex mixtures^2^. A common targeted proteomics technique is the use of a triple quadrupole MS to collect selected reaction monitoring (SRM) data. The number of analytes that can be measured in triple quadrupole assays is limited. Thus, following digestion of a protein mixture, precursor and product ion pairs (transitions) are selected from peptides to provide a proxy measurement of the protein of interest. The high selectivity and sensitivity of SRM results in accurate and reproducible peptide quantification^3^. It is possible to validate the measurement of each analyte through figures of merit including limit of detection and quantification, ensuring that we can produce robust and reliable results^4^. However, the primary challenge in establishing a targeted assay is selecting peptides that are 1) good proxies of abundance for specific proteoforms or combinations of proteoforms^5^, 2) respond well in the mass spectrometer from the target sample matrix^6–8^, and 3) chromatograph well^9,10^.

Common peptide selection techniques rely on manual or automated analysis of empirical data found in literature review, databases of targeted measures, and discovery experiments^1,10–13^. Confounding factors for peptide and transition selection for SRM include differences in peptide response due to sample matrix, sample preparation, separation, and analysis instrumentation and methods. For example, the majority of public data from discovery experiments was collected by data dependent acquisition (DDA), and these data are often leveraged to select proxy peptides or train predictive algorithms. However, the criteria that result in frequent sampling and confident identification of DDA spectra (e.g. abundant MS1 signal, many characteristic fragments, wide elution peaks) are specifically poor performers for a targeted analysis (e.g., a symmetrical and narrow chromatographic peak, selective precursor > product ion pairs of high abundance)^9,10,14^. An alternative strategy to DDA for evaluating peptide suitability is implementing SRM to assess recombinant proteins of interest. Full or near-full length protein is purified and digested, then peptides are measured individually to determine target transitions for a quantitative assay^9^. Although this approach eliminates the issue of translating DDA performance to SRM experiments, it remains a challenge to determine performance of endogenous peptides in the specific matrix of interest. Thus, the selection of peptides for SRM is often a laborious and expensive process, requiring the purchase or recombinant production of a protein standard, multiple rounds of data acquisition and peptide target filtering^1,9,14,15^.

Data-independent acquisition (DIA) is an alternative MS acquisition strategy that measures the proteome in a more systematic manner. Predictive methods trained on DIA data have been shown to outperform DDA-based approaches for predicting which peptides will perform well in a targeted assay^14^. Additionally some have used DIA data to aid in the development of parallel reaction monitoring methods^16,17^. We previously reported a method that used multiple injections to cover the mass range for typical tryptic peptides using DIA with narrow isolation windows^18^ that is analogous to the PAcIFIC method ^19,20^. These narrow, effectively 2 *m/z*, isolation windows across the relevant mass range, are similar in size to parallel reaction monitoring (PRM) methods ^21^. Data can be collected directly from biological samples of interest, thus taking into account the influence of the sample matrix. Here we leverage a gas phase fractionated library collected from biological samples to efficiently select peptides for SRM. The indexed retention time (iRT) values and transitions are inherently determined during the generation of the library, making SRM scheduling and transition selection seamless ^22^. The workflow minimizes the number of comprehensive measurements to assess which peptides 1) are good responders in the matrix of interest, 2) chromatograph well, and 3) exhibit stability and precision for measurement. To demonstrate the usefulness of this workflow we compare peptides selected from narrow-window DIA to those included in an established, well-characterized SRM assay for Alzheimer’s disease in cerebrospinal fluid for a set of proteins ^1^. Using the workflow presented herein, an assay for the same proteins of interest was developed, exhibiting equivalent precision and accuracy but bypassing multiple rounds of selection and refinement. Additionally, the technique described is implemented to generate multiple separate quantitative assays for different targets from the same initial data.

## METHODS

### Cerebrospinal fluid patient samples

Cerebrospinal fluid (CSF) samples were obtained by lumbar tap from patients diagnosed as either probable Alzheimer’s disease or Parkinson’s disease, and healthy age-matched controls as a part of the Alzheimer’s disease resource center at the University of Washington, or the Udall repository at Stanford University. CSF was obtained following NIA AD Center Best Practices Guidelines and samples stored at −80°C. Equal volumes of 30 Parkinson’s disease, 11 probable Alzheimer’s disease, and 30 non-disease CSF samples were combined to make a pooled reference for method development. All samples were pre-existing and were collected under protocols approved by the Institutional Review Board (IRB) at University of Washington or Stanford University prior to the start of this project. The UW and Stanford Human Subjects Division deems the use of pre-existing de-identified samples exempt from full IRB review and, thus, treated this project as non-human subjects research.

### Cerebrospinal fluid sample preparation

CSF was diluted 1:1 with 0.1% PPS silent surfactant (Expedeon) in 100 mM ammonium bicarbonate with 3.5 ng/μl of [15N]APOA1, and heated to 95°C for 5 minutes to facilitate protein denaturation. All samples were reduced with 5 mM dithiothreitol, alkylated with 15 mM iodoacetamide, and the alkylation reaction quenched with 5 mM dithiothreitol. Proteins were digested with sequencing grade trypsin (Pierce) at a 1:25 enzyme to substrate ratio for 16 h at 37°C. Reactions were quenched and PPS surfactant hydrolyzed with 6N hydrochloric acid to reach pH 2. Samples were desalted by solid phase extraction (SPE) using Waters Oasis 60 μm/30 mg MCX cartridges (Milford, MA). The SPE resin was washed and conditioned with 1 mL of methanol, 1 mL of 2.8% ammonium hydroxide in water, 2 mL of methanol, and 3 mL of 0.1% formic acid (FA) in water. Equilibration of the SPE resin was performed using 1 mL of 0.1% FA in water, and 0.1% FA in methanol. Acidified digested samples were loaded onto SPE resin, washed with 1 ml 0.1% FA in water and 1 ml of 90% acetonitrile/10% water, and peptides eluted with 1 mL of 2.8% ammonium hydroxide in methanol. Peptides were dried by vacuum centrifugation and stored at −80°C. Samples were resuspended to 0.33 μg/μl in 0.1% FA in water with Pierce retention time calibrator (PRTC) peptides spiked into each sample to a final concentration of 15 mM.

### DIA-MS and processing

Tryptic peptides were separated using a Thermo EASY nanoLC 1200 on self-packed 30 cm columns packed with 3 μm ReproSil-Pur C18 silica beads (Dr. Maisch, Ammerbuch, Germany) in a 75 μm inner diameter fused silica capillary (#PF360 Self-Pack PicoFrit, New Objective). Trap columns were created from 150 μm inner diameter fused silica capillary fritted with Kasil on one end and packed with the same C18 beads to 25 mm. Solvent A was 0.1% formic acid in water and solvent B was 0.1% formic acid in 98% acetonitrile. For each injection, 1 μg of peptide was loaded and eluted using a 90 min gradient from 5 to 35% B, followed by a 40 min washing gradient. The 30 cm column was heated to 35°C. Data were acquired by narrow window DIA experiments with a Thermo Q-Exactive HF tandem mass spectrometer, as described in^18,23^. For each DIA experiment 6 GPF acquisitions were performed with 4 *m/z* precursor isolation windows at 30,000 resolution, 1e6 target AGC, 55 ms maximum injection times, and NCE of 27. Narrow mass range windows were staggered with optimized window placements as detailed in Pino et al.^23^ Two MS1 scan events were collected every 25 MS2 spectra. One of the MS1 scan events spanned the 400-1000 m/z range covered by all gas phase fractionated runs. The second MS1 scan event covered just the 100 m/z range covered by the specific MS2 isolation windows using the selected ion monitoring (SIM) scan function to improve the MS1 dynamic range. DIA data were demultiplexed with a 10 ppm tolerance following peak picking by ProteoWizard MSConvert ^24^. Using Walnut, a re-implementation of PECAN^25^, available through EncyclopeDIA (version - 0.6.8), resulting mzMLs were searched using with a Uniprot reviewed human proteome (downloaded 6/2016) with APOE variant peptides appended, configured with the default settings: fixed cysteine carbamidomethylation, 10 ppm precursor and product tolerances, using y-ions, and assuming full tryptic digestion with up to 1 missed cleavage. EncyclopeDIA search results were then filtered to a 1% peptide-level FDR using Percolator 3.1 and then filtered again to a 1% protein-level FDR.

### Selected reaction monitoring acquisition and processing

Targeted data were acquired using a SRM-MS method on a Thermo TSQ Altis triple-quadrupole instrument using a Thermo EASY nLC 1200 for separation. Approximately 1 μg of each sample was separated on self-packed 15 cm columns packed with 3 μm ReproSil-Pur C18 silica beads (Dr. Maisch, Ammerbuch, Germany) in a 75 μm inner diameter fused silica capillary (#PF360 Self-Pack PicoFrit, New Objective). Peptides were separated using a 60 min gradient from (2% acetonitrile in 0.1% formic acid to 23% acetonitrile in 0.1% formic acid). The gradient was followed by a wash for 10 min at 80% acetonitrile in 0.1% formic acid and a column re-equilibration at 0.1% formic acid for 10 min. Ions were isolated in both Q1 and Q3 using 0.7 FWHM resolution and peptide fragmentation was performed at 1.5 mTorr in Q2 using calculated peptide specific collision energies. Data was acquired using a scan width of 0.002 mass to charge ratio (*m/z*) and a dwell time of 10 ms, for 5 minutes around the predicted retention time. Targeted proteomic data were analyzed using the software package Skyline^26,27^. Chromatographic peak intensities from all monitored transitions of a given peptide were integrated and summed to give a final peptide peak area. TIC normalization, statistical analysis, and figure generation were performed with R.

## Data availability

The Skyline documents, raw files, DIA data, and supplementary data are available at Panorama Public^28^. All code used for the data normalization and figure generation is available online at https://github.com/uw-maccosslab/manuscript-dia-to-srm.

## RESULTS

### DIA provides a list of sample specific detectable peptides

A total of 18 ug of CSF digest was injected across 18 injections for the 3 replicate, 6 gas-phase fractionated (GPF) DIA libraries. From the three GPF-DIA libraries, 6661 peptides corresponding to 1176 proteins were detected across all replicates (Supplemental Figure 1). 5595 of the detected peptides, corresponding to 1154 proteins, were determined to be possible SRM targets based on the criteria of a minimum of 3 co-eluting transitions with no interference. Interference was determined by EncyclopeDIA as transitions with a peak apex not co-eluting with the other transitions. For all possible targets, precursor retention times and relative product ion intensities were extracted in Skyline for quantification. A percent coefficient of variation (%CV) was calculated for every peptide to determine signal stability across 3 days in a 4°C autosampler. For all possible target peptides the median %CV was 13.5%, with 4089 peptides corresponding to 1018 proteins having %CV ≤ 20 (Supplemental Figure 2). Typically, peptides that have more product ions with no interference have more reproducible abundances (Supplemental Figure 3).

### Using peptide characteristics from DIA measurements to filter out poor responding peptides

Using the peptide %CV calculations and information about product ion transition intensities, peptides were selected that are expected to perform well in a triple-quadrupole targeted experiment. Since the %CV is calculated from triplicate DIA experiments over separate days, the %CV captures peptide stability in an autosampler at 4 °C. For the 4564 peptides with calculated %CV less than 20, transitions were ranked by measured intensity for each peptide and peptide summed area for the top 5 interference-free transitions was used to rank peptides mapping to the same target protein. To build the SRM assays (Figure 1) the selection criteria were the following; (1) for proteins with more than 2 peptides only the top 5 peptides, as ranked by signal intensity, were considered. (2) From the top 5 ranked peptides, the 2 peptides with the lowest %CV were selected to be included in our final assay. (3) For the 2 peptides with the lowest %CV, the top 5 most intense product ion transitions were included in the targeted assay.

**Figure 1.**
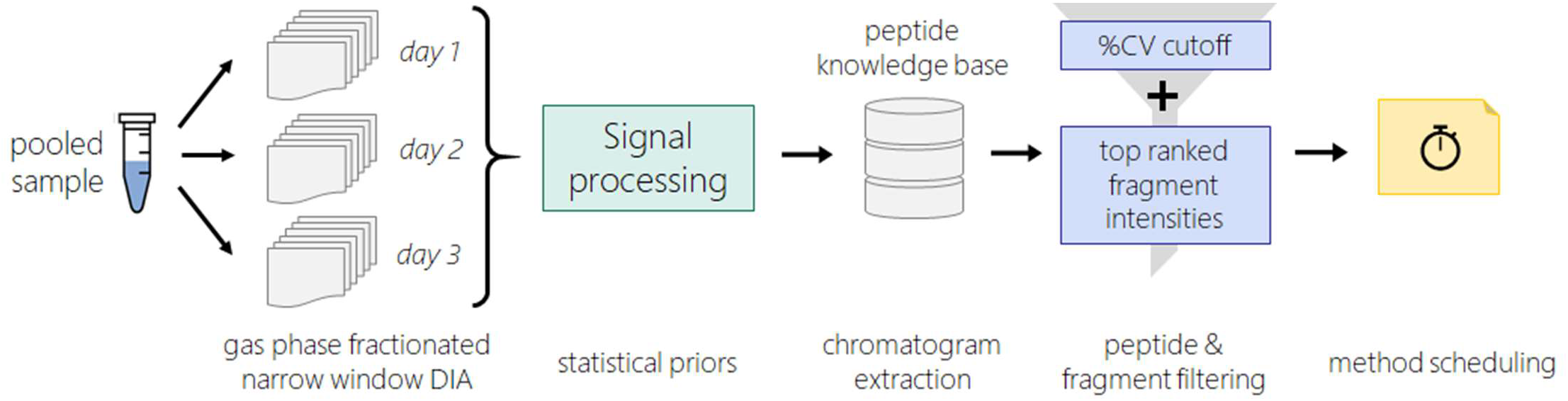
Workflow for selecting peptides and transitions for targeted assay. A pooled sample of cerebrospinal fluid from a neurodegenerative disease cohort was subjected to 3 gas-phase fractionated data independent acquisition experiments. Each gas-phase fractionated DIA experiment required 6 injections, and spanned the 400-1000 *m/z* range in 100 *m/z* increments. Peptide detection, relative retention time, and precursor product ion pairs are determined through a peptide-centric search method. Search results and chromatographic information is extracted in Skyline to form a knowledge base used for quantitation and visualization. For peptides with 3+ transitions that co-elute and have no interference a %CV is calculated between the summed peptide area across the 3 DIA experiments. For proteins that have more than 2 peptides with %CV less than 30, peptides are ranked by summed area for up to 5 transitions. For the top ranking peptides the 5 most intense transitions are selected for inclusion in targeted assay list.

### Performance of peptides for previously targeted Alzheimer’s disease proteins

To test the robustness of peptide selection for SRM assays based on DIA data, a targeted assay was developed based on a well characterized assay for use in Alzheimer’s dementia research (Spellman). From the 180 proteins targeted in a published assay ^1^ peptides mapping to 170 were detected by GPF-DIA in a CSF pool, and 96 of these proteins were detected with at least 2 unique peptides. For those 96 proteins we selected up to two of the previously targeted peptides to measure by SRM. From the GPF-DIA data we also selected up to 2 peptides to target based on the aforementioned criteria. This resulted in three proteins (HBA, APP, VASN) with the same two peptides selected from both, and three protein isoforms with the same unique peptides targeted (NEO1, APOE-E2, APOE-E4). 51 proteins had no overlap in the peptides selected from either data. Approximately 74% of the selected peptides from the GPF-DIA were uniquely selected, with about 26% (50 peptides) overlapping with the previously published targeted assay (Figure 2). Peptides and transitions selected from both the previous assay and from the GPF-DIA data were used to create a single inclusion list. Scheduled windows of 5 minutes and limiting the number of concurrent transitions to 200 resulted in 4 separate methods with transitions from the 13 PRTC peptides included in each method. For the peptides selected from DIA the performance is comparable with the previous assay, with a median CV of 3.6%, and 3.2% respectively (Figure 2).

**Figure 2.**
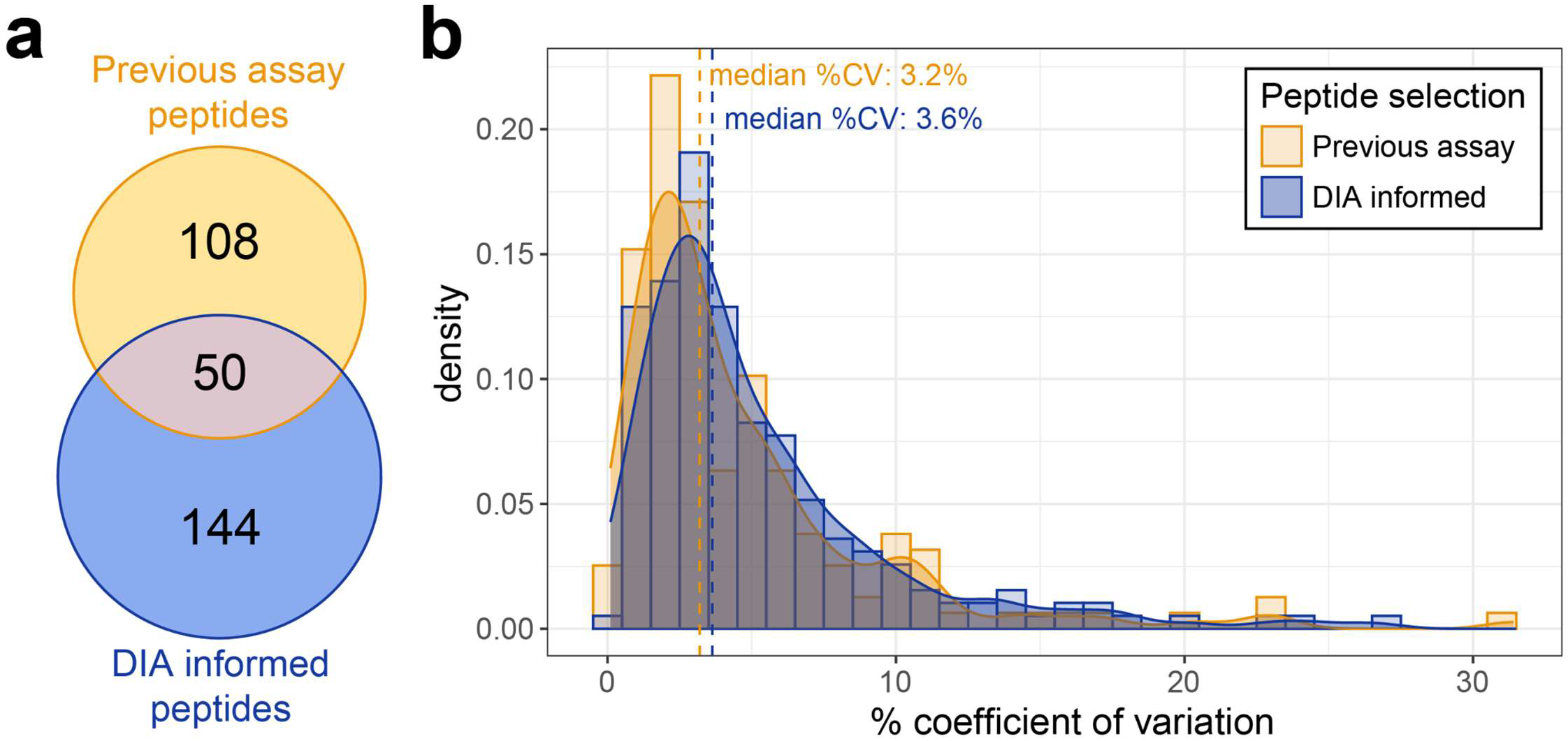
Peptides selected from narrow window DIA results perform similarly to previously characterized Alzheimer’s disease assay peptide selections. For 98 proteins previously included in a targeted Alzheimer’s disease assay, we targeted peptides either selected by the previous study (yellow) or selected using data from our DIA experiments (blue). We measured 5 transitions for peptides previously targeted, and from our selection. The percent coefficient of variation is calculated from 3 replicate injections of pooled cerebrospinal fluid measured by SRM and plotted by selection process.

### DIA data can be re-queried to select peptides for additional protein assays

One benefit of acquiring DIA data to inform assay generation is that we obtain valuable peptide information for all detected peptides. This means the data can be used to generate multiple different targeted assays depending on the proteins of interest in any given biological question or application. To demonstrate our ability to generate a separate assay for a different subset of proteins from a GPF-DIA dataset, the same CSF peptide knowledge base data was used to develop a second SRM assay for proteins previously described as being differential in CSF sampled from sufferers of chronic pain. From the 87 proteins found to be differential in the Khoonsari et al. study ^29^, peptides were detected for 85 proteins by GPF-DIA. Of the detected proteins, 67 were detected by at least 2 peptides with at least 5 co-eluting and interference-free transitions. The two best performing peptides per protein based on %CV and relative intensity ranking were included in the targeted assay, resulting in 134 target peptides and 670 target transitions. A median %CV of 6.2 across 3 replicate acquisitions of pooled CSF by the chronic pain SRM assay (Figure 3).

**Figure 3.**
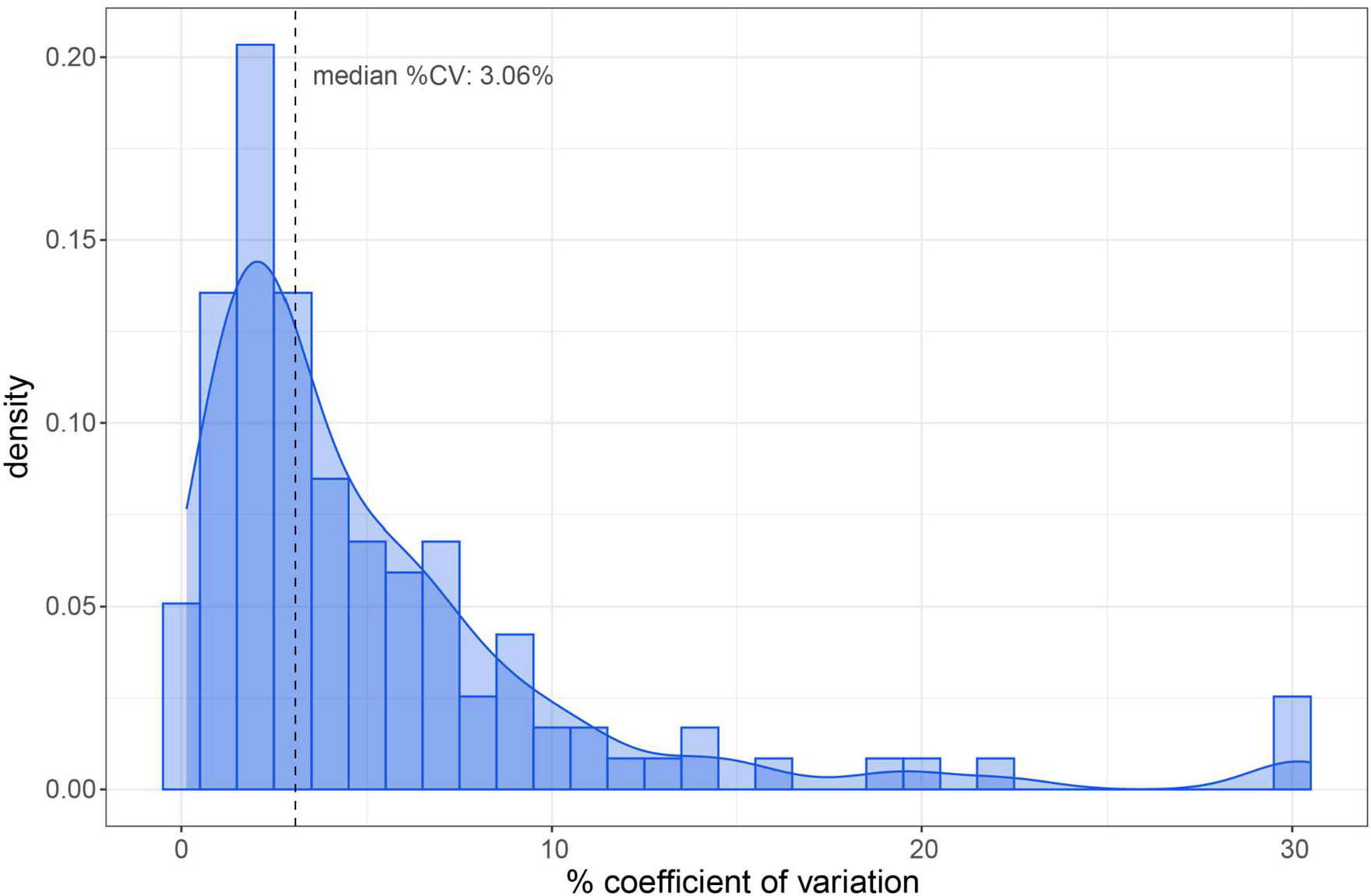
Additional assays generated using the same DIA experiments. Peptides for targeted triple quadrupole measurement were selected for 67 previously described pain proteins using information about transition signal and chromatographic performance from DIA. For the 134 peptides measured in triplicate the median %CV of 3.06.

## DISCUSSION

Here we demonstrate a workflow to generate a triple quadrupole SRM assay with just a couple days of instrument time, minimal technician time, little reagent cost, and relatively small amounts of sample (less than 10 ug of protein). Peptides and transitions with high intensity, good chromatographic performance, and good precision by GPF-DIA were selected for inclusion. The utility of this GPF-DIA to SRM workflow was demonstrated by reproducing a well characterized, previously published SRM assay for AD based on the DIA data. The re-use of the GPF-DIA was demonstrated through the generation of a separate reproducible assay for the quantification of proteins that are of interest as markers of chronic pain.

This workflow is especially efficient because DIA data is collected using a representative, pooled sample of interest. From this data the chromatographic performance of each detectable peptide in the biological matrix of interest can be rapidly evaluated, which is valuable information for our selection of targets. By sampling the 400-1000 *m/z* range with 2 *m/z* windows we are able to determine which peptide species are detectable in our sample type of interest with the ability to distinguish between different sequences with similar precursor masses^18,25^. The inclusion of indexed retention time standards in the sample analyzed by DIA allows for transfer of relative retention time between different liquid chromatography setups and facilitates scheduling of a SRM assay if necessary^22^. This step also highlights the importance of measuring peptides in the background matrix of interest for the final targeted assay because peptide retention time and intensity is dependent on the matrix^30,31^.

The use of DIA methods has increased in recent years, leading to the accumulation of a plethora of data that could be used for the development of targeted assays. These existing datasets can often be implemented to provide information regarding which peptides are likely to be of interest for a targeted reanalysis of these proteins. Narrow window DIA can be acquired as “chromatogram” libraries as part of larger DIA studies as described by Searle et al. and Pino et al. to improve the analysis of the data^18,23^. This workflow provides quantitative data for comparisons in discovery cohorts, which can further inform the selection of targets. We show that the information gained from one set of DIA experiments can be mined for multiple purposes, making DIA data a valuable resource for future studies. Previous work from Pino et al demonstrates the use of DIA for more extensive characterization of peptide quantification through matrix matched calibration curves^6^. This could be included as an additional step in determining suitable targets for SRM assay development. Further considerations could also be made to optimize the selection of peptides to maximize the number of precursors across a gradient, by selecting peptides that distribute concurrent peptides across time.

Triple quadrupole assays are a valuable method for the rapid and robust, targeted analysis of select analytes. The selection of peptides and target transitions for analysis has remained a time and labor intensive bottleneck in assay development. This step often makes use of prior DDA information followed by multiple rounds of validation experiments to determine which peptides perform well by SRM, or it relies on previous SRM assays which will be limited in measured sample types or targets. The workflow presented here greatly reduces the upfront time and effort required to generate a well-performing assay. By acquiring replicate gasphase fractionated DIA and refining targets in the Skyline environment, we were able to pick target peptides and transitions with good intensity, chromatographic performance, and precision in a triple quad assay with only several days of instrument and data processing time.

## ACKNOWLEDGEMENTS

This work was supported in part by the National Institute of Health grants U19 AG065156, R24 GM141156, F31 AG069420, and U01 DK137097. Cerebrospinal fluid was provided by the Adult Changes in Thought study (R13 AG057087), and Stanford Udall Center (P50 NS062684).

