## Supplementary Figures for "Data Independent Acquisition to Inform the Development of Targeted Proteomics Assays Using a Triple Quadrupole Mass Spectrometer"

**SUPPLEMENTAL FIGURES.**

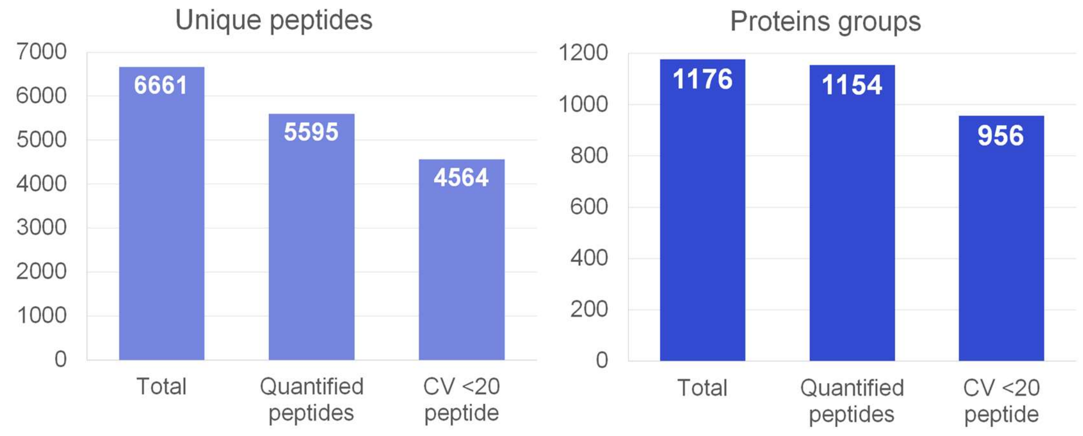


**Supplemental Figure 1. Number of protein and peptide detections by DIA experiments.** The total number of detections from 3 replicate experiments is plotted for both proteins and peptides (Total). The number of proteins with at least one peptide with at least 3 interference-free product ion transitions (Quantified Peptides). Peptides were filtered for those that exhibit a peak area CV of less than 20% over the 3 GPF libraries (CV <20 peptide).

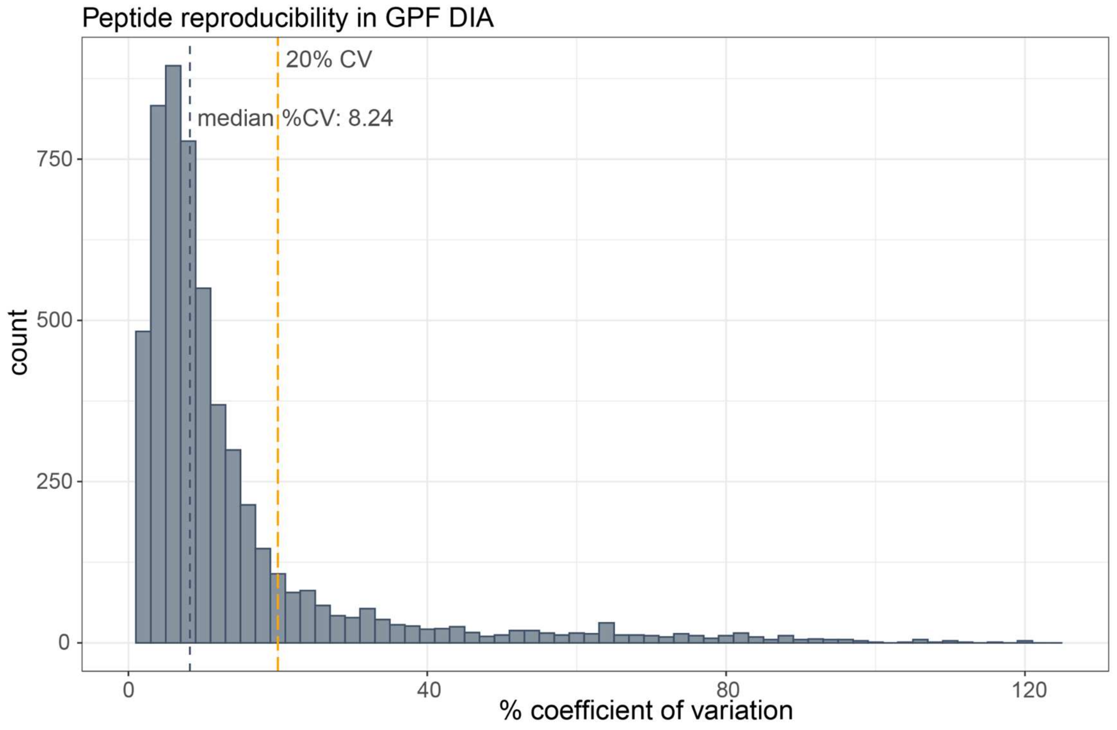


**Supplemental Figure 2. Peptide reproducibility across 3 gas-phase fractionated DIA experiments.** The percent coefficient of variation for all peptides detected by DIA with at least 3 interference-free transitions. The median CV is 8.24%, below the typical threshold of 20% as indicated by the dashed line.

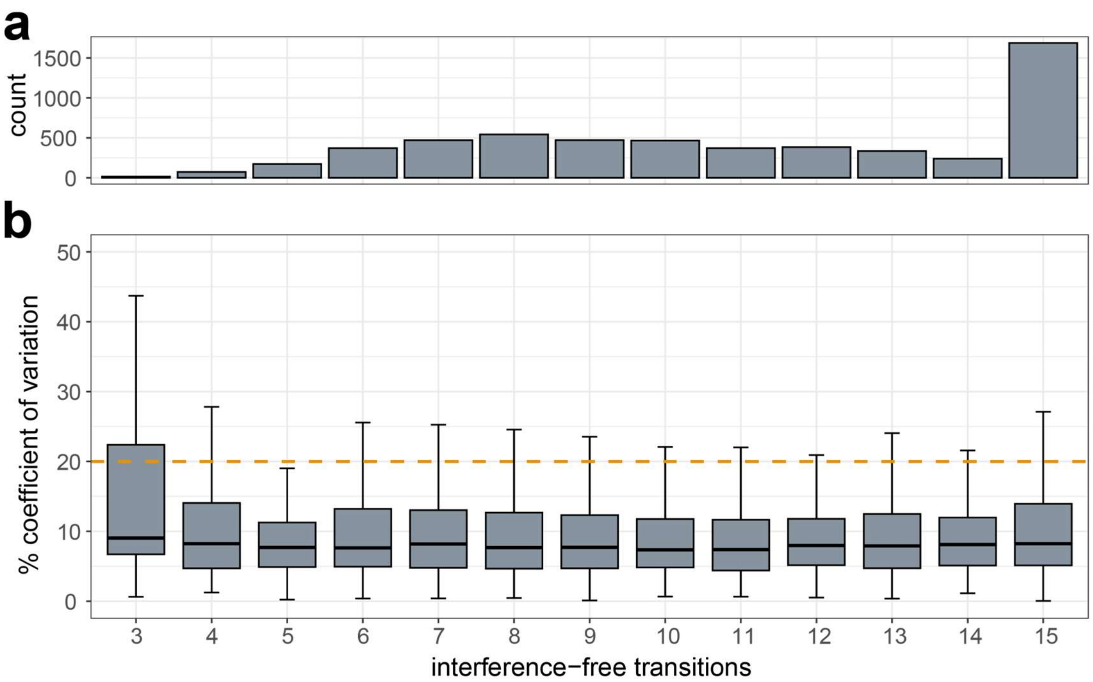


**Supplemental Figure 3. Reproducibility is improved with the number of product ion transitions detected per peptide.** Using EncyclopeDIA we determine which transitions are interference-free. For peptides that have at least 3 interference-free transitions we extract data in Skyline for quantitation. a) The number of interference-free product ions for each peptide detected by GPF-DIA. The majority (%) of peptides quantified contain more than 5 product transitions. b) The percent coefficient of variation (%CV) from 3 replicate gas-phase fractionated narrow window DIA experiments. The median %CV decreases with an increased number of interference-free transitions per peptide.

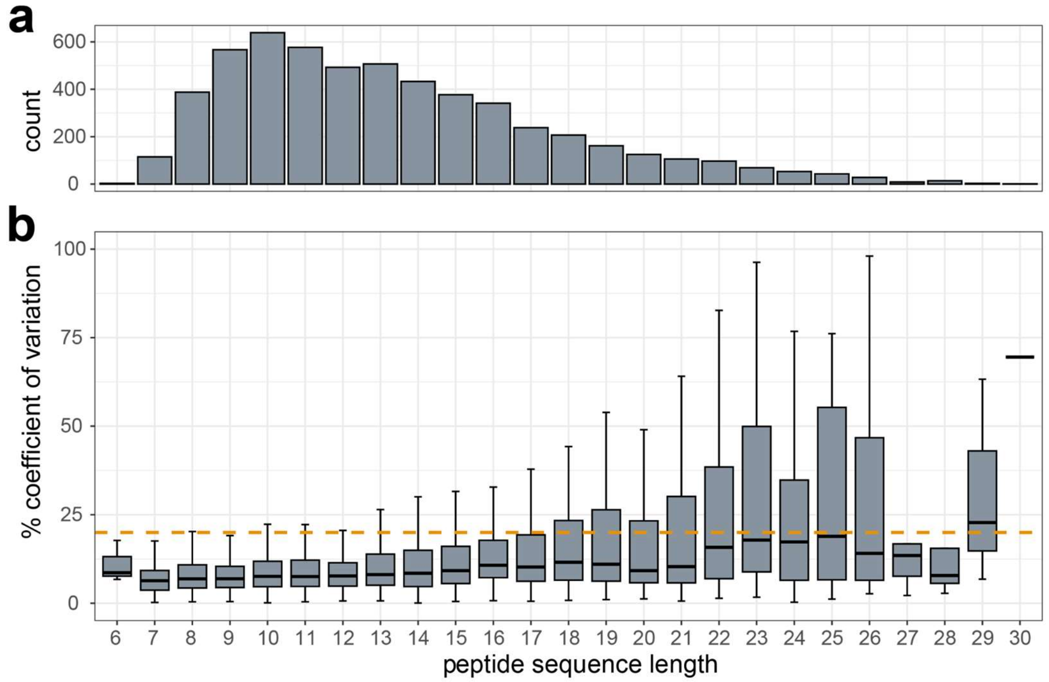


**Supplemental Figure 4. Reproducibility of peptides decreases with increasing sequence length.** a) The distribution of peptides found in the GPF-DIA by their peptide sequence length. b) The percent coefficient of variation (%CV) from 3 replicate gas-phase fractionated narrow window DIA experiments. The median %CV increases as sequence length increases.
